# Fecal metabolomics reveals preferential complex carbohydrate utilization and guides cultivation of murine gut *Firmicutes*

**DOI:** 10.64898/2026.08.19.745854

**Authors:** Preethi Sudhakara, James P. Martin, Joan A. Whitlock, Timothy J. Garrett, Gurjit S. Sidhu, Gary P. Wang

**Affiliations:** Division of Infectious Disease and Global Medicine, Department of Medicine, University of Florida, Gainesville, FL, USA; Department of Pathology, Immunology and Laboratory Medicine, College of Medicine, University of Florida, Gainesville, FL, USA; Infectious Diseases Section, Medical Service, North Florida/South Georgia Veterans Health System, Gainesville, FL, USA

**Keywords:** murine gut microbiota, *Firmicutes* cultivation, novel media, fecal metabolomics, Clostridioides difficile, colonization resistance, carbon catabolite repression, PYE6S

## Abstract

The murine gut microbiota provides robust colonization resistance against *Clostridioides difficile* infection (CDI), yet murine-associated microbes remain notoriously difficult to cultivate in vitro, limiting mechanistic investigation. To identify the ecological and nutritional basis of this cultivation barrier, we leveraged CDI susceptibility as a functional readout of microbial community metabolism to infer *in vivo* nutrient utilization. Germ-free C57BL/6 mice colonized with varying dilutions of ethanol-treated murine microbiota were challenged with *C. difficile* resulting in a spectrum of CDI outcomes. Comparative metabolomics of pre-challenge fecal samples revealed a consistent carbohydrate signature: glucose accumulated in communities that resisted *C. difficile* challenge, whereas complex carbohydrates, including raffinose, sucrose, trehalose, lactose, sorbitol, and mannitol, were significantly depleted. The broad depletion of these complex carbohydrates supports their functional importance within the collective microbial community. Conventional glucose-based media (CMA, BHI+I, RCMT) failed to support robust growth or subculture of murine gut microbiota. Guided by the metabolomics findings, we developed Peptone Yeast Extract with Six Salts and Sugars (PYE6S), a glucose-free medium supplemented with the complex carbohydrates identified as depleted. PYE6S enabled cultivation of 22 unique *Firmicutes* ASVs, 82% of which lacked named cultured representatives in reference databases. These findings suggest a plausible explanation for why conventional media fail and support a metabolomics-guided framework for rational cultivation of host-associated microbiota across diverse systems. This strategy may be extended to guide media design for other host-associated microbiotas.

## Introduction

The gastrointestinal tract harbors a complex microbial community that plays a central role in colonization resistance against enteric pathogens^1,2^. This function is particularly important in *Clostridioides difficile* infection (CDI), one of the leading causes of healthcare-associated diarrhea, responsible for approximately half a million cases annually in the United States^3,4^. Fecal microbiota transplantation achieves sustained clinical cure in over 90% of patients with recurrent CDI when administered repeatedly^5,6^, demonstrating the importance of the gut microbiota in mediating resistance. Members of the *Firmicutes* phylum (*Bacillota*), particularly Lachnospiraceae and Ruminococcaceae, are consistently depleted in CDI and have been proposed as key contributors to colonization resistance^7–9^.

Mouse models have been indispensable for studying microbiota-mediated CDI resistance. Germ-free mice are uniformly lethal when challenged with *C. difficile*, whereas colonization with murine microbiota restores protection, implicating specific microbial communities in CDI resistance^10^. However, a major limitation remains: murine gut microbes are notoriously difficult to cultivate using conventional laboratory media, preventing isolation and functional investigation of individual species. Recent efforts such as the Mouse Intestinal Bacterial Collection (miBC) and Mouse Gut Bacterial Collection (MGBC) have demonstrated that cultivation is possible^11,12^. However, a mechanistic understanding of why standard media fail, and how to rationally design media that support cultivation of fastidious organisms, remains incomplete.

Conventional anaerobic media use glucose as the primary carbon source. While this approach supports cultivation of many laboratory-adapted organisms, it may be poorly suited to gut *Firmicutes* that have adapted to a colonic environment in which free glucose is largely absent, as glucose is absorbed in the proximal small intestine. Colonic substrates consist predominantly of complex polysaccharides, host-derived glycans, and microbially derived metabolites. In addition, many gut *Firmicutes* are inefficient metabolizers of free glucose despite encoding the genetic machinery to sense it^13–15^. Furthermore, in Gram-positive bacteria, glucose activates carbon catabolite repression (CCR) via formation of an HPr-Ser-P–CcpA complex that binds catabolite-responsive elements (cre) and represses genes involved in complex carbohydrate utilization^16,17^. We hypothesized that conventional glucose-based media fail to support cultivation of murine gut *Firmicutes* via two potential mechanisms. First, glucose is a poor nutrient substrate to support sustained growth of these organisms. Second, glucose concurrently engages CcpA-mediated repression of the polysaccharide utilization pathways these organisms require. Thus, we reasoned that defining the metabolomic environment of the murine cecum would identify the substrates required for sustained growth in vitro, thereby guiding the rational design of media that recapitulate the colonic substrate environment.

To test this hypothesis, we used a gnotobiotic model in which germ-free mice were colonized with increasing dilutions of ethanol-treated murine microbiota, generating a spectrum of communities with differing metabolic capacity. We used CDI susceptibility as a functional readout to stratify communities by metabolic state. By comparing pre-challenge fecal metabolomes across these states, we aimed to deduce *in vivo* nutrient utilization preferences of murine gut microbes that could guide rational medium design.

This approach revealed a consistent carbohydrate signature distinguishing protective from non-protective communities against *C. difficile*, which directly informed the design of a novel glucose-free culture medium. Here, we describe the metabolomic insights and their application to overcoming the cultivation barrier, providing a generalizable framework for microbiome cultivation based on *in vivo* metabolic preferences. This strategy offers a broadly applicable approach for inferring nutrient utilization and guiding the cultivation of host-associated microbiota across diverse systems.

## Material and Methods

### Animals and *C. difficile* challenge

All procedures were approved by the University of Florida Institutional Animal Care and Use Committee and the University of Florida Institutional Review Board. Fecal pellets were collected from wild-type C57BL/6 conventional mice and humanized C57BL/6 mice previously colonized with human stools from another study. A 10% (w/v) fecal suspension was prepared in pre-reduced 1× phosphate-buffered saline (PBS, pH 7.2), centrifuged at 10,000 × g for 10 minutes, and the pellet was treated with 70% ethanol for 2 hours to eliminate vegetative cells. The ethanol-resistant fraction was then centrifuged, washed twice with PBS, and resuspended for oral gavage.

Germ-free (GF) C57BL/6 mice were orally gavaged with the ethanol-resistant fraction and individually housed in HEPA-filtered NexGen cages with autoclaved food and water in a dedicated ABSL2 suite. Fecal pellets were then collected, and a 10% (w/v) suspension prepared in PBS. This suspension was serially diluted (1:10, 1:100, 1:1000), and 100 µL of each dilution was gavaged into GF C57BL/6 mice (n = 29 total; 1:10 dilution, n = 15; 1:100, n = 7; 1:1000, n = 1; undiluted positive controls, n = 3; PBS negative controls, n = 3). After 14 days of colonization, fecal samples were collected for metabolomics and 16S rRNA sequencing. Mice were then challenged with 300-400 CFU of *C. difficile* VPI 10463 spores by oral gavage and monitored daily for clinical signs including diarrhea and weight loss for up to 14 days. Mice exhibiting severe diarrhea and/or weight loss exceeding 15% of pre-challenge body weight were euthanized according to humane endpoint criteria.

For *C. difficile* enumeration, cecal contents were suspended in 1× PBS (10% w/v), serially diluted, and plated on taurocholate cycloserine cefoxitin fructose agar (TCCFA)^18^ under anaerobic conditions at 37°C. Colonies were enumerated after 48 hours of incubation.

### Fecal metabolomics

Fecal samples collected prior to *C. difficile* challenge were subjected to metabolite extraction using a cellular extraction protocol with normalization to total protein content. Global untargeted metabolomics profiling was performed using a Thermo Q-Exactive Orbitrap mass spectrometer coupled with a Dionex UHPLC system and autosampler. Samples were analyzed in both positive and negative heated electrospray ionization modes in separate injections, with a mass resolution of 35,000 at m/z 200. Chromatographic separation was achieved using an ACE 18-pfp column (100 × 2.1 mm, 2 µm particle size) with mobile phase A consisting of 0.1% formic acid in water and mobile phase B consisting of acetonitrile, at a flow rate of 350 µL/min and column temperature of 25°C. Injection volumes were 4 µL (negative mode) and 2 µL (positive mode).

### Comparative fecal metabolomics analysis

A total of 7,526 features were detected in positive ion mode and 6,754 features in negative ion mode. Data were normalized to the total ion signal per sample. Feature detection, alignment, deisotoping, and gap filling were performed using MZmine^19^. Adducts and complex ions were removed, and metabolite identification was performed using an internal retention time library. Statistical analysis was performed separately for positive and negative ion datasets, comparing samples from mice that survived *C. difficile* challenge versus those that did not. Univariate analysis was conducted using volcano plots, with a significance threshold of p < 0.05 applied to known metabolites. Given the exploratory nature of this analysis, unadjusted p-values were used to prioritize biologically interpretable metabolites. The full list of significantly altered metabolites is provided in Table S1.

### 16S rRNA gene sequencing and bioinformatics

DNA was extracted from fecal samples and individual isolates using the Chelex method. A 10% Chelex solution was prepared in Tris-EDTA (TE) buffer, approximately 1 µL of each isolate was resuspended in the Chelex solution, mixed by gentle tapping, and heated at 100°C for 15 minutes. Following lysis, samples were centrifuged at 10,000 × g for 2 minutes and the supernatant transferred to TE buffer and stored at −20°C until sequencing.

The V1–V3 hypervariable region of the 16S rRNA gene was amplified using forward primer 27F (5′-AGA GTT TGA TCC TGG CTC AG-3′) and reverse primer 534R (5′-ATT ACC GCG GCT GCT GG-3′), incorporating barcodes for multiplex sequencing as described previously^20^. The final 20 µL PCR reaction contained 0.75 U Accuprime Taq High Fidelity Polymerase (Invitrogen, Carlsbad, CA), 2 µL 10× PCR buffer II, 2 µM forward primer, 2 µM reverse primer, and 2 µL DNA template. Thermal cycling conditions were as follows: initial denaturation at 95°C for 2 minutes, followed by 25 cycles of denaturation at 95°C for 20 seconds, annealing at 56°C for 30 seconds, and extension at 72°C for 5 minutes. PCR products were analyzed on a 1% SYBR Safe agarose gel, and amplicons of the expected size (~500 bp) were excised and purified using the NucleoSpin Gel and PCR Clean-up kit (Macherey-Nagel, Bethlehem, PA). DNA concentration was quantified using the Qubit HS DNA quantification kit (Invitrogen, Carlsbad, CA). Equimolar concentrations were pooled and library concentration verified by qPCR using the Library Quant Kit (Kapa Biosystems, Wilmington, MA). A 6 pM library was sequenced on an Illumina MiSeq platform using the MiSeq Reagent Kit v3.

MiSeq reads were demultiplexed using cutadapt^21^, imported into QIIME 2 v2024.10^22^, and denoised using the DADA2 pipeline. Taxonomy was assigned using the classify-sklearn algorithm against the SILVA 138 reference database, and manuscript nomenclature follows SILVA assignments where applicable. Organism names used for GenBank sequence deposition follow the current NCBI Taxonomy nomenclature, which may differ from the SILVA taxonomy for recently reclassified taxa. A Maximum Likelihood phylogenetic tree based on 16S rRNA gene sequences of the cultured isolates was constructed using MEGA12^23^. Statistical analyses and visualization were performed using GraphPad Prism v10.1.

### Microbial growth on culture media

Ethanol-treated fecal suspensions (10% w/v) derived from conventional mice or humanized mice were plated on PYE6S, Reinforced Clostridial Medium with taurocholate (RCMT), Cooked Meat Agar (CMA), and Brain Heart Infusion supplemented with inulin (BHI+I, 0.1% inulin). Plates were incubated anaerobically at 37°C. Colonies from primary plates were transferred to secondary plates of the same medium to assess sustainability of colony growth. Only isolates with robust growth on both primary and secondary plates were selected for downstream analysis.

### Statistical analysis

Group comparisons were performed using the Mann–Whitney test (two-group comparisons). Volcano plot analysis for metabolomics data used a significance threshold of p < 0.05. All data are presented as group means ± standard deviation (SD). Statistical analyses were performed using GraphPad Prism v10.1. A p-value < 0.05 was considered statistically significant.

## Results

### Conventional culture media fail to support robust growth of murine gut microbiota

We first examined whether conventional anaerobic culture media could support growth of gut microbes from the ethanol-treated fraction of murine gut microbiota, and compared this with growth from ethanol-treated humanized mouse fecal microbiota under identical conditions (Figure 1). Fecal suspensions from humanized germ-free mice colonized with ethanol-treated human microbiota yielded abundant colonies on Cooked Meat Agar (CMA) and Brain Heart Infusion supplemented with inulin (BHI+I), with visible growth within 2–3 days and robust subculture on secondary plates. In contrast, ethanol-treated murine fecal suspensions plated on CMA, BHI+I, and Reinforced Clostridial Medium with taurocholate (RCMT) produced only sparse, small colonies after prolonged incubation (5–11 days). Importantly, when these colonies were transferred to secondary plates of the same media, the vast majority of murine microbiota failed to grow, suggesting that initial colony formation was likely supported by residual nutrients present in the fecal inoculum rather than by the culture media itself. This observation of delayed primary growth followed by failure of subculture suggests that conventional glucose-based media do not provide the substrates required for sustained growth of murine gut microbes, and indicate that murine-associated *Firmicutes* may have distinct nutritional requirements not met by standard laboratory media.

**Figure 1.**
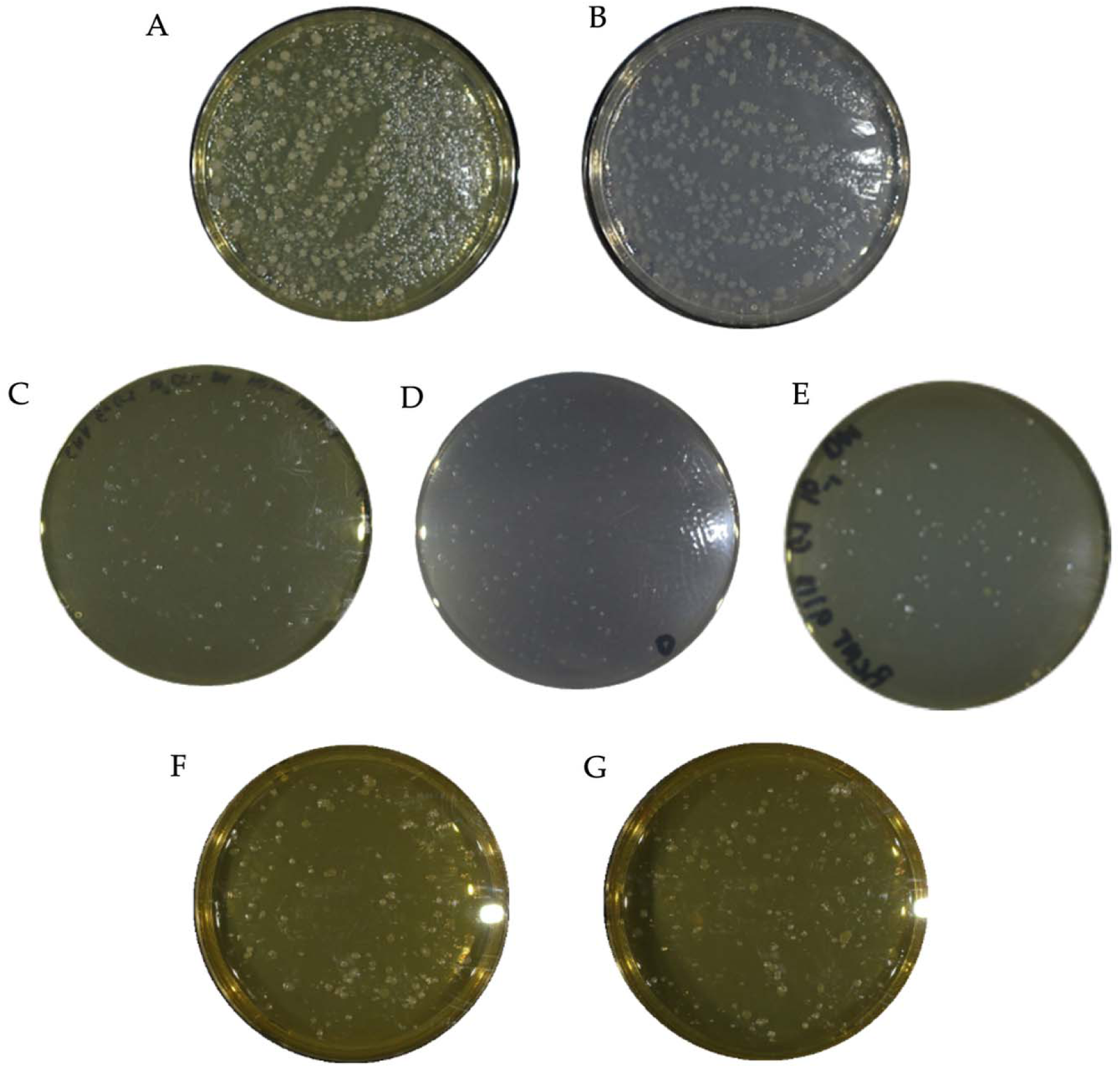
Conventional media fail, while PYE6S robustly supports cultivation of murine gut microbes enriched for *Firmicutes*. Colonies recovered from fecal suspensions plated under anaerobic conditions on conventional media or the metabolomics-guided PYE6S medium. (A,B) Ethanol-treated humanized mouse fecal suspension plated on Cooked Meat Agar (CMA) and Brain Heart Infusion supplemented with 0.1% inulin (BHI+I), respectively, showing abundant colony growth by day 3. (C–E) Ethanol-treated conventional mouse fecal suspension plated on CMA (day 5), BHI+I (day 7), and Reinforced Clostridial Medium with taurocholate (RCMT; day 10), demonstrating sparse delayed colony formation that failed to subculture. (F,G) Ethanol-treated conventional mouse fecal suspension plated on Peptone Yeast Extract with Six Salts and Sugars (PYE6S), showing abundant colony growth by day 6 on two replicate plates. These findings demonstrate that conventional glucose-based media poorly support cultivation of murine gut *Firmicutes*, whereas the glucose-free PYE6S formulation enables robust growth.

### Serial dilution of murine microbiota produces a spectrum of CDI susceptibility

To determine the *in vivo* metabolic requirements of murine gut *Firmicutes*, we established a gnotobiotic model in which serial dilutions of ethanol-treated murine microbiota generated communities with varying levels of microbial diversity and metabolic capacity (Figure 2). Germ-free C57BL/6 mice (n = 29) colonized with serial dilutions of the ethanol-resistant fraction were challenged with *C. difficile* VPI 10463 following 14 days of colonization. Mice colonized with undiluted ethanol-resistant fraction (positive controls, n = 3) were resistant to *C. difficile* challenge with no detectable cecal *C. difficile*, and maintained stable body weight and clinical health scores throughout the 14-day observation period (Figure S1). In contrast, germ-free negative controls rapidly succumbed to *C. difficile* challenge, reaching humane endpoints within 36 hours post-challenge.

**Figure 2.**
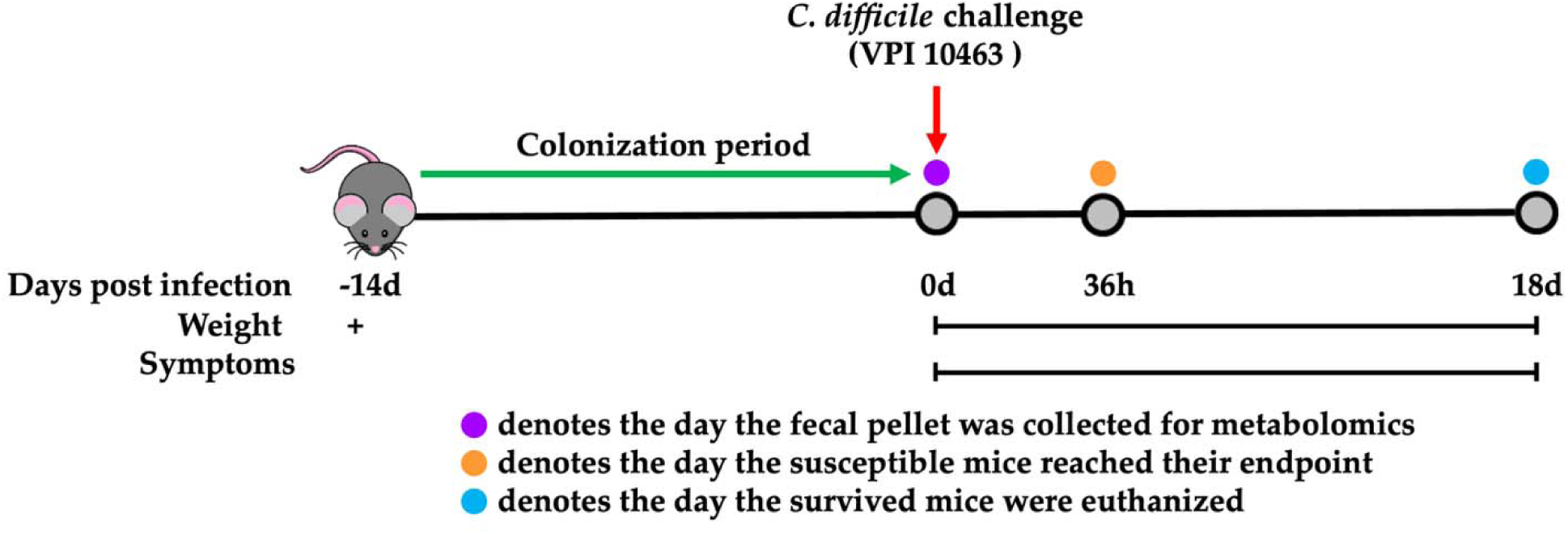
Experimental design. Schematic of the *C. difficile* infection and metabolomics study. Germ-free (GF) C57BL/6 mice were colonized with serially diluted ethanol-treated murine fecal microbiota for 14 days. Fecal samples were collected prior to *C. difficile* challenge for metabolomics and 16S rRNA gene sequencing. Mice were then challenged orally with 300-400 CFU of *C. difficile* VPI 10463 spores and monitored for 14 days.

Among the mice colonized with diluted microbiota, three distinct CDI phenotypes were observed following *C. difficile* challenge. Resistant mice (n = 5) remained clinically healthy throughout the 14-day observation period, had no detectable *C. difficile* in the cecum, and were negative for TcdA/B by qualitative assay. Symptomatic mice (n = 7) survived *C. difficile* challenge but had diarrhea and were persistently positive for *C. difficile* and TcdA/B. Susceptible mice (n = 17) developed severe diarrhea with weight loss exceeding 15% of pre-challenge body weight, and were toxin-positive with high cecal *C. difficile* burdens at the time of euthanasia. Disease severity correlated with dilution level, with higher dilutions associated with increased susceptibility.

### Comparative fecal metabolomics reveals preferential utilization of complex carbohydrates

To identify substrates preferentially utilized by murine microbiota *in vivo*, we performed comparative metabolomic analysis of pre-challenge fecal samples from mice that survived versus those that did not survive *C. difficile* challenge. Across all samples, 7,526 features were detected in positive ion mode and 6,754 features in negative ion mode. Univariate volcano plot analysis identified a consistent carbohydrate signature distinguishing between the two groups. Specifically, raffinose, trehalose, sucrose, lactose, sorbitol, and mannitol were significantly depleted in mice that survived *C. difficile* challenge, whereas glucose/fructose was significantly enriched (p < 0.05 for all; Figure 3, Table S1), indicating a depletion of multiple complex carbohydrates with concurrent accumulation of glucose. These results are consistent with preferential utilization of complex carbohydrates *in vivo* by murine gut communities associated with *C. difficile* protection. While substrate utilization by individual taxa was not specifically examined, the broad depletion of multiple structurally distinct, complex carbohydrates supports the hypothesis that complex carbohydrates are preferentially utilized by the murine *Firmicutes*. The observed accumulation of glucose may reflect reduced utilization, or production from polysaccharide metabolism, or a combination of both. Taken together, these findings indicate that the nutritional landscape of the murine gut differs substantially from that of conventional culture media, which typically include glucose as the primary carbon source.

**Figure 3.**
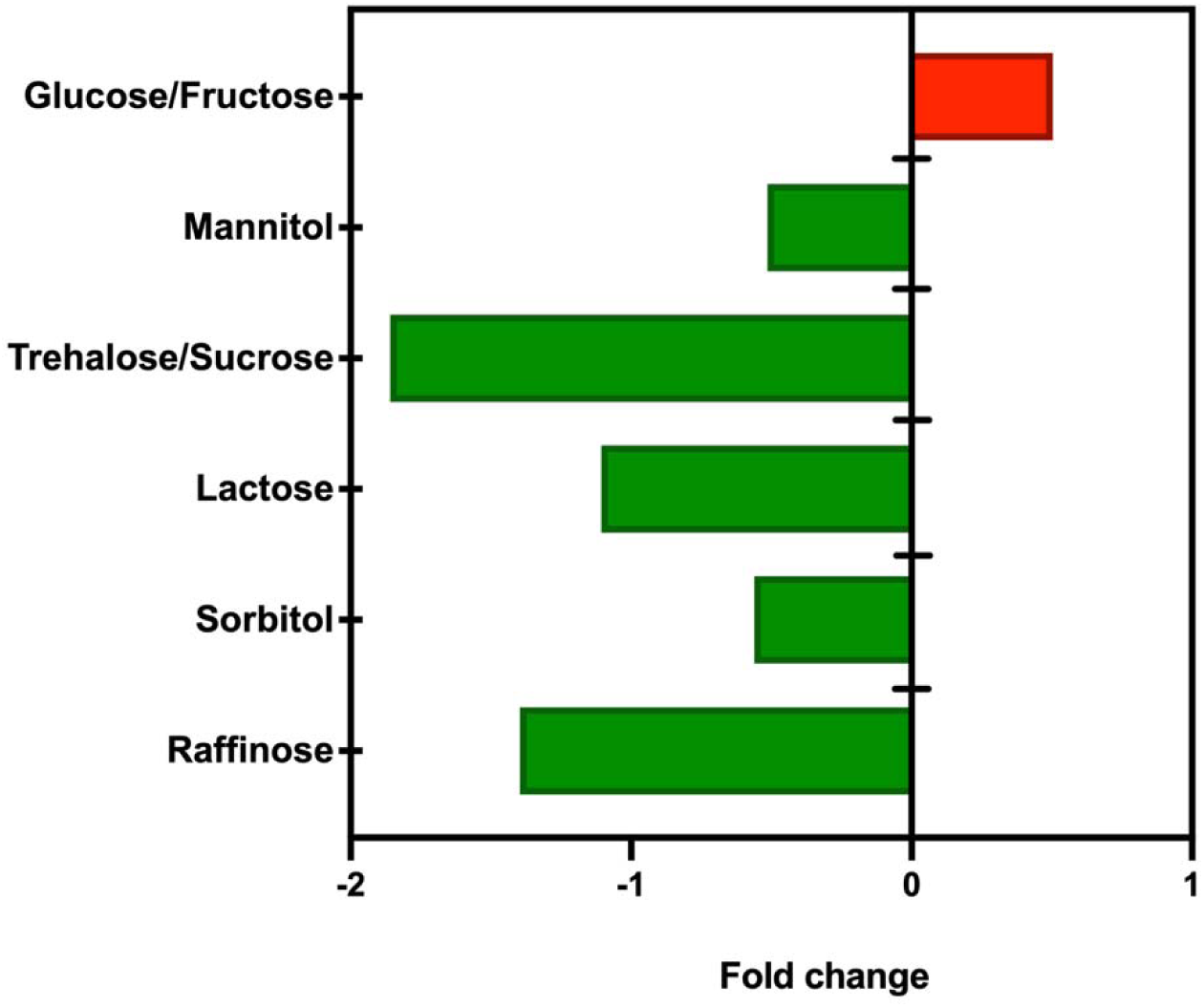
Comparative fecal metabolomics reveals preferential utilization of complex carbohydrates. Fold changes of significantly altered sugars in the pre-challenge fecal metabolome of mice that survived versus mice that died following *C. difficile* challenge (p < 0.05, volcano plot analysis). Green bars indicate metabolites depleted in survivors, consistent with microbial consumption. Red bars indicate metabolites enriched in survivors. Raffinose, trehalose/sucrose, lactose, sorbitol, and mannitol were significantly depleted, whereas glucose/fructose was significantly enriched.

### Rational design of a metabolomics-guided culture medium

Informed by the metabolomics findings, we designed a culture medium that replaces glucose with the complex carbohydrates identified as depleted *in vivo*. Sucrose, trehalose, lactose, mannitol, sorbitol, and raffinose were incorporated as components of the carbohydrate fraction. Pancreatic digest of casein (Casitone) and yeast extract were used as nitrogen source in place of animal-derived extracts present in CMA and BHI to better reflect the murine colonic environment adapted to typical plant-based murine diet. An inorganic salt solution was included, and medium pH was empirically optimized. pH 6.8 yielded the highest colony recovery (87 colonies), compared with pH 6 (1 colony), pH 7.5 (3 colonies), and pH 8 (1 colony). pH 6.8 falls within the range of measured murine cecal and colonic pH values^24^, which span approximately pH 5.5–6.7 depending on diet and microbiota composition. The resulting formulation, Peptone Yeast Extract with Six Salts and Sugars (PYE6S), is detailed in Table 1.

**Table 1.** Composition of Peptone Yeast Extract with Six Salts and Sugars (PYE6S)

| **Component** | **Amount per 1 L** |
| --- | --- |
| Pancreatic Digest of Casein | 20 g |
| Yeast Extract | 10 g |
| L-Cysteine Hydrochloride | 0.5 g |
| Agar | 15 g |
| DI Water | 860 mL |
| Autoclave; then add: |  |
| *25X Salts Solution | 40 mL |
| #10% Sugars Solution | 100 mL |
| *25X Salts Solution (per 100 mL) |  |
| Calcium Chloride (anhydrous) | 0.02 g |
| Magnesium Sulfate | 0.02 g |
| Potassium Phosphate Monobasic | 0.1 g |
| Potassium Phosphate Dibasic | 0.1 g |
| Sodium Chloride | 0.2 g |
| Sodium Bicarbonate | 1 g |
| Dissolve in 100 mL DI water; filter sterilize |  |
| #10% Sugars Solution (per 100 mL) |  |
| Sucrose | 1.6 g |
| Trehalose | 1.6 g |
| Lactose | 1.6 g |
| Mannitol | 1.6 g |
| Sorbitol | 1.6 g |
| Raffinose | 1.6 g |
| Dissolve in 90 mL DI water; filter sterilize |  |
| Add sugar and salt solutions in anaerobic chamber after autoclave. Adjust to pH 6.8. |  |

### PYE6S supports robust cultivation and subculture of murine *Firmicutes*

In contrast to the sparse and delayed growth observed on conventional media, PYE6S yielded robust growth of colonies derived from ethanol-treated murine fecal suspensions by day 6 (Figure 1). A total of 87 colonies were picked from primary plates, and all (100%) were successfully sub-cultured on secondary PYE6S plates.

16S rRNA sequencing of the 87 purified colonies identified 22 unique Amplicon Sequence Variants (ASVs), all belonging to the *Firmicutes* phylum (Table 2). Of these, 18 (82%) lacked named species-level cultured representatives in the reference database, whereas 4 (18%) matched previously described taxa. Phylogenetic analysis revealed three major clades (Figure 4). Clade 1 comprised members of Ruminococcaceae and Oscillospiraceae, including *Clostridium leptum* and *Acutalibacter muris*. Clade 2 included *Lachnoclostridium* and related taxa within Lachnospiraceae. Clade 3 encompassed members of Erysipelotrichaceae and related lineages within Peptostreptococcales and Erysipelotrichales. At the family level, isolates were distributed across Lachnospiraceae (36.4%), Oscillospiraceae (22.7%), Ruminococcaceae (18.2%), Erysipelotrichaceae (9.1%), Clostridiaceae (4.6%), Anaerovoracaceae (4.6%), and Erysipeloclostridiaceae (4.6%) (Figure S2). Because all isolates were derived from ethanol-treated fecal samples, these organisms are inferred to be spore-formers.

**Figure 4.**
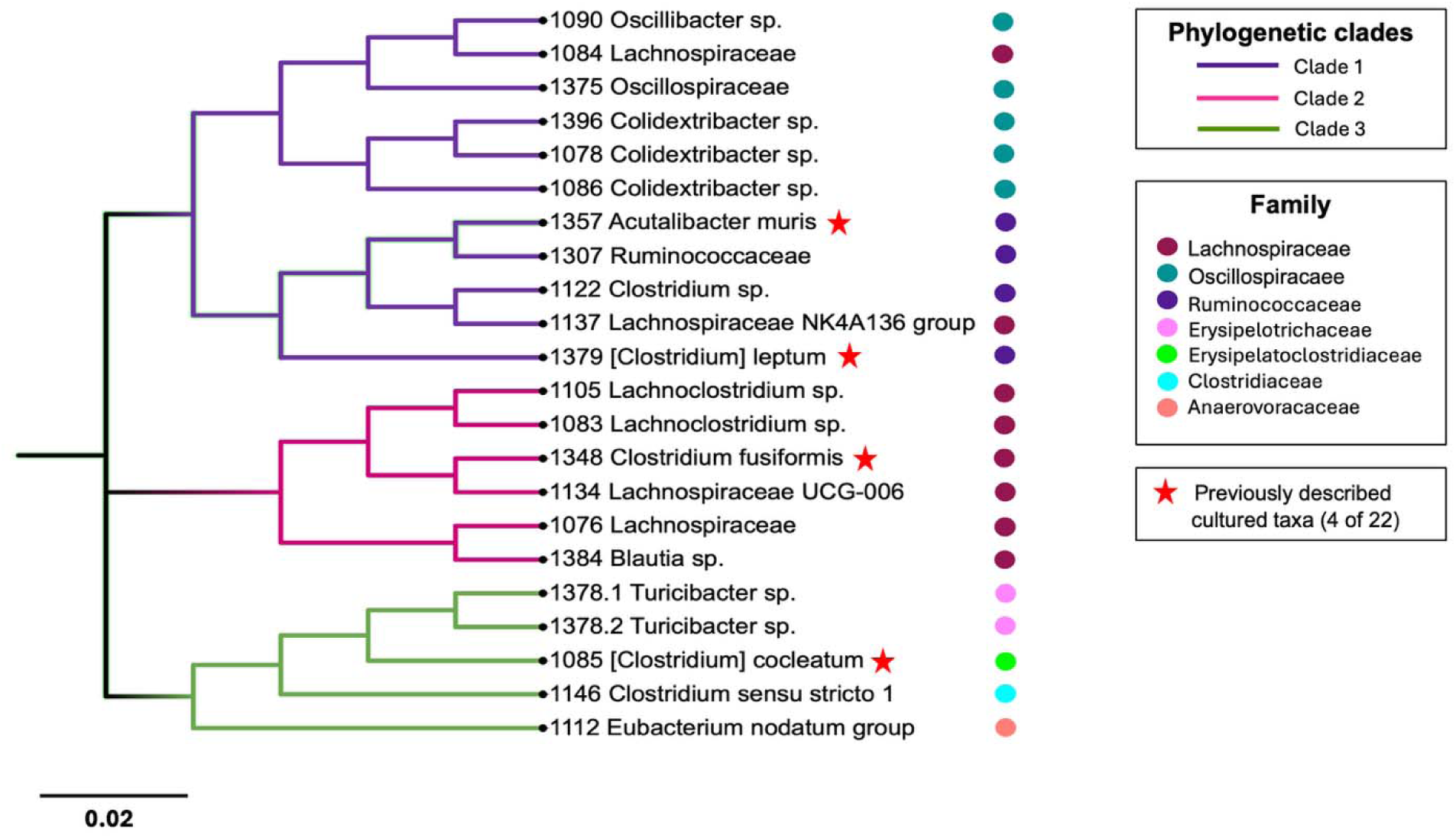
Phylogenetic analysis of 22 murine *Firmicutes* isolates cultivated on PYE6S. Maximum-likelihood phylogenetic tree based on V1–V3 16S rRNA gene sequences of the 22 cultivated ASVs. Three major phylogenetic clades are highlighted in purple, pink, and green. Colored dots indicate family-level taxonomic assignments based on the SILVA 138 reference database. Red stars indicate the four previously described cultured taxa. The scale bar represents 0.02 substitutions per site. Eighteen of the 22 isolates lacked named cultured representatives in the SILVA 138 reference database, whereas four corresponded to previously described cultured taxa.

**Table 2.** Taxonomy of the 22 ASVs cultivated using PYE6S. The table includes 18 novel cultured bacterial isolates and 4 previously described cultured bacterial taxa. Eighteen ASVs lacked named cultured representatives in the reference database, whereas four matched previously described cultured taxa. Taxonomic assignments were generated using the SILVA 138 reference database with QIIME 2. Bracketed taxon names ([Clostridium]) indicate historical genus names retained in the SILVA taxonomy following taxonomic reclassification. Corresponding GenBank accession numbers are provided for each isolate. Organisms’ names used for GenBank sequence deposition follow the current NCBI Taxonomy nomenclature.

| ASVs | Isolate ID | Class | Order | Family | Genus | Species | Accession Number |
| --- | --- | --- | --- | --- | --- | --- | --- |
| **Novel cultured bacterial isolates** |  |  |  |  |  |  |  |
| 1 | 1076 | Clostridia | Lachnospirales | Lachnospiraceae | *Unclassified* | *Unclassified* | PZ639008 |
| 2 | 1105 | Clostridia | Lachnospirales | Lachnospiraceae | *Lachnoclostridium* | *Unclassified* | PZ639009 |
| 3 | 1090 | Clostridia | Oscillospirales | Oscillospiraceae | *Oscillibacter* | *Unclassified* | PZ639010 |
| 4 | 1112 | Clostridia | Peptostreptococcales-Tissierellales | Anaerovoracaceae | [*Eubacterium*] *nodatum* group | *Unclassified* | PZ639012 |
| 5 | 1122 | Clostridia | Oscillospirales | Ruminococcaceae | *Incertae Sedis* | *Clostridium* *species* | PZ639013 |
| 6 | 1375 | Clostridia | Oscillospirales | Oscillospiraceae | *Unclassified* | *Unclassified* | PZ639014 |
| 7 | 1146 | Clostridia | Clostridiales | Clostridiaceae | *Clostridium sensu stricto 1* | *Unclassified* | PZ639015 |
| 8 | 1307 | Clostridia | Oscillospirales | Ruminococcaceae | *Incertae Sedis* | *Unclassified* | PZ639016 |
| 9 | 1396 | Clostridia | Oscillospirales | Oscillospiraceae | *Colidextribacter* | *Unclassified* | PZ639017 |
| 10 | 1086 | Clostridia | Oscillospirales | Oscillospiraceae | *Colidextribacter* | *Unclassified* | PZ639018 |
| 11 | 1384 | Clostridia | Lachnospirales | Lachnospiraceae | *Blautia* | *Lachnospiraceae* *bacterium* | PZ639019 |
| 12 | 1078 | Clostridia | Oscillospirales | Oscillospiraceae | *Colidextribacter* | *Unclassified* | PZ639027 |
| 13 | 1083 | Clostridia | Lachnospirales | Lachnospiraceae | *Lachnoclostridium* | *Unclassified* | PZ639021 |
| 14 | 1084 | Clostridia | Lachnospirales | Lachnospiraceae | *Uncultured* | *Unclassified* | PZ639022 |
| 15 | 1137 | Clostridia | Lachnospirales | Lachnospiraceae | *Lachnospiraceae_*<br>*NK4A136_group* | *Unclassified* | PZ639025 |
| 16 | 1134 | Clostridia | Lachnospirales | Lachnospiraceae | *Lachnospiraceae_*<br>*UCG-006* | *Clostridium*<br>*species* | PZ639026 |
| 17 | 1378.1 | Bacilli | Erysipelotrichales | Erysipelotrichaceae | *Turicibacter* | *Turicibacter*<br>*species* | PZ639028 |
| 18 | 1378.2 | Bacilli | Erysipelotrichales | Erysipelotrichaceae | *Turicibacter* | *Unclassified* | PZ639029 |
| ***Previously described Cultured bacteria*** |  |  |  |  |  |  |  |
| 19 | 1085 | Bacilli | Erysipelotrichales | Erysipelatoclostridiaceae | *Erysipelatoclostridium* | *[Clostridium]*<br>*cocleatum* | PZ639020 |
| 20 | 1348 | Clostridia | Lachnospirales | Lachnospiraceae | *Lachnoclostridium* | *Clostridium*<br>*fusiformis* | PZ639023 |
| 21 | 1379 | Clostridia | Oscillospirales | Ruminococcaceae | *Anaerotruncus* | *[Clostridium]*<br>*leptum* | PZ639024 |
| 22 | 1357 | Clostridia | Oscillospirales | Ruminococcaceae | *Incertae Sedis* | *Acutalibacter*<br>*muris* | PZ639011 |

## Discussion

This study addresses a longstanding and poorly understood limitation in murine microbiome research—the inability of conventional media to support cultivation of many murine gut microbes. By integrating comparative fecal metabolomics with a gnotobiotic model that generated three distinct CDI phenotypes (resistant, symptomatic carrier, and susceptible), we identified an *in vivo* metabolic signature that directly informed an in vitro cultivation strategy. These findings link community metabolic state to culturability and offer a rational framework to overcome a significant experimental barrier in murine microbiome research.

A central finding of this study was the broad depletion of multiple complex carbohydrates—including raffinose, trehalose, sucrose, lactose, sorbitol, and mannitol—in microbial communities associated with survival following *C. difficile* challenge, accompanied by relative accumulation of glucose. The *in vivo* depletion of structurally distinct carbohydrates aligns with active microbial utilization, suggesting murine gut *Firmicutes* are adapted to a metabolic environment where complex dietary or host-derived carbohydrates, not free glucose, are primary carbon sources.

The failure of conventional glucose-based media to support sustained growth of murine gut *Firmicutes* may reflect two complementary mechanisms. First, these organisms are adapted to a colonic environment in which free glucose is largely absent because glucose is absorbed in the proximal small intestine, and instead rely on complex polysaccharides, host-derived glycans, and microbially derived metabolites for growth^13–15^. Second, glucose is known to activate CcpA-mediated carbon catabolite repression (CCR) in Gram-positive bacteria through formation of an HPr-Ser-P–CcpA complex that binds catabolite-responsive elements (cre) and represses genes involved in utilization of alternative carbohydrates^16,17^. Thus, glucose-based media may be suboptimal for growth, concurrently suppressing genes crucial for metabolizing more physiologically relevant substrates. This interpretation may also explain the delayed primary growth and failure of subculture observed on conventional media, where residual nutrients carried over in the fecal inoculum may transiently support initial colony formation but are depleted upon subculture. Although we did not directly measure CcpA activity, transcriptional responses, or glucose uptake, the *in vivo* depletion of complex carbohydrates with concurrent enrichment of glucose, together with the inability of these organisms to sustain growth in glucose-based media, is consistent with this mechanistic interpretation.

This interpretation may also help explain the success of prior cultivation efforts. Large-scale murine microbiota collections such as miBC^11^ and MGBC^12^ recovered diverse murine commensals using media formulations containing complex carbohydrates, mucin, rumen fluid, or related substrates, with or without glucose. Such formulations may provide carbon sources that these organisms can utilize efficiently while reducing dependence on free glucose as the dominant substrate. Our findings offer a plausible explanation for why these approaches were effective and suggest that in vitro cultivation may be improved by more closely recapitulating the *in vivo* nutrient environment. A conceptual advance of the present study is the use of CDI susceptibility as a functional readout to stratify communities by metabolic activity, enabling inference of nutrient utilization without requiring prior knowledge of the nutritional requirements of individual taxa.

The PYE6S medium developed here supported robust growth and subculture of murine *Firmicutes* and recovered substantial taxonomic diversity, including a high proportion of taxa lacking named cultured representatives in reference databases. The cultivated organisms were notably enriched for families such as Lachnospiraceae, Ruminococcaceae, and Oscillospiraceae, all associated with colonization resistance in murine and human studies^7,8,10^. Although the present study does not establish functional roles for these isolates, their recovery provides a foundation for future mechanistic studies using defined strains or consortia. The substrate-mismatch hypothesis proposed here may also extend to nutritionally fastidious human gut *Firmicutes*, including *Faecalibacterium prausnitzii*, which is known to depend on complex carbohydrates, host-derived glycans, and cross-fed metabolites for sustained growth^13^.

This study has several limitations. First, the metabolomic analyses of fecal samples do not allow assignment of specific metabolic activities to individual taxa. Second, although the proposed CCR-based mechanism provides a plausible explanation for the observed cultivation barrier, transcriptional responses, glucose uptake, and direct CcpA activity were not examined and remain important areas for future study. Third, while the concurrent depletion of multiple complex carbohydrates with relative enrichment of glucose is consistent with microbial liberation of simple sugars from polysaccharides, reduced glucose consumption may also contribute to this observation. Finally, we did not directly compare PYE6S with previously published murine isolation media under identical conditions, and such comparison would be informative in future studies.

Our findings suggest that the longstanding challenge of cultivating gut *Firmicutes* reflects, at least in part, a mismatch between conventional laboratory media and the *in vivo* nutrient environment, rather than intrinsic uncultivability. More broadly, comparative metabolomics across distinct community states may provide a generalizable strategy for identifying substrates that support growth in vitro. By aligning culture conditions more closely with the metabolic environment determined *in vivo*, such approaches may help expand the cultivable diversity of host-associated microbial communities and facilitate future mechanistic studies of previously uncultivable organisms.

## Acknowledgements

We thank members of the Wang laboratory for helpful discussions.

## Author Contributions

Study concept and design: G.P.W. and G.S.S. Acquisition of data: P.S., J.P.M., J.A.W. Metabolomics: G.S.S. and T.J.G. Analysis and interpretation of data: P.S., G.S.S., and G.P.W. Drafting of manuscript: P.S., G.S.S., and G.P.W. Critical revision: P.S., G.S.S., G.P.W.

## Data Availability

Demultiplexed FASTA sequences of the cultured isolates are available in the DANS Data Station Life Sciences repository at https://doi.org/10.17026/LS/VV66OE, DANS Data Station Life Sciences, V2. 16S rRNA gene sequences of the 22 isolates: NCBI GenBank [PZ639008–PZ639029].

