## Supplemental figures and table for "Fecal metabolomics reveals preferential complex carbohydrate utilization and guides cultivation of murine gut *Firmicutes*"

**Supplemental Material:**

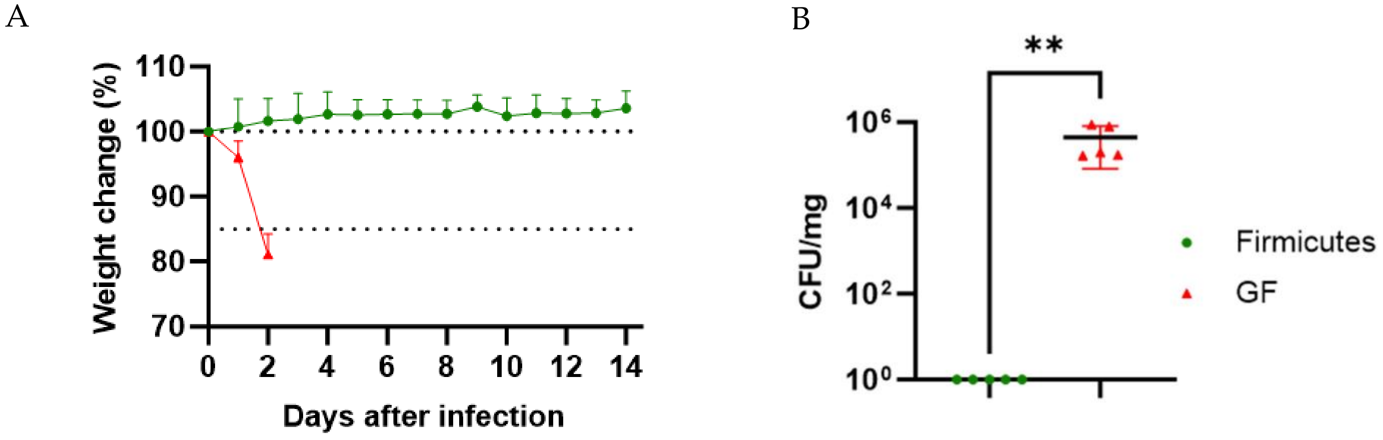


**Figure S1. *Firmicutes*-colonized mice resist *C. difficile* challenge.** (A) Body weight change (%) over 14 days post-infection in *Firmicutes*-colonized mice (green) and GF negative controls (red). GF mice reached humane endpoints by day 2. (B) Cecal *C. difficile* burden at endpoint: *Firmicutes*-colonized mice at day 14 (below detection) versus GF mice at euthanasia (>$10^5$ CFU/mg). **P < 0.05, Mann–Whitney test.

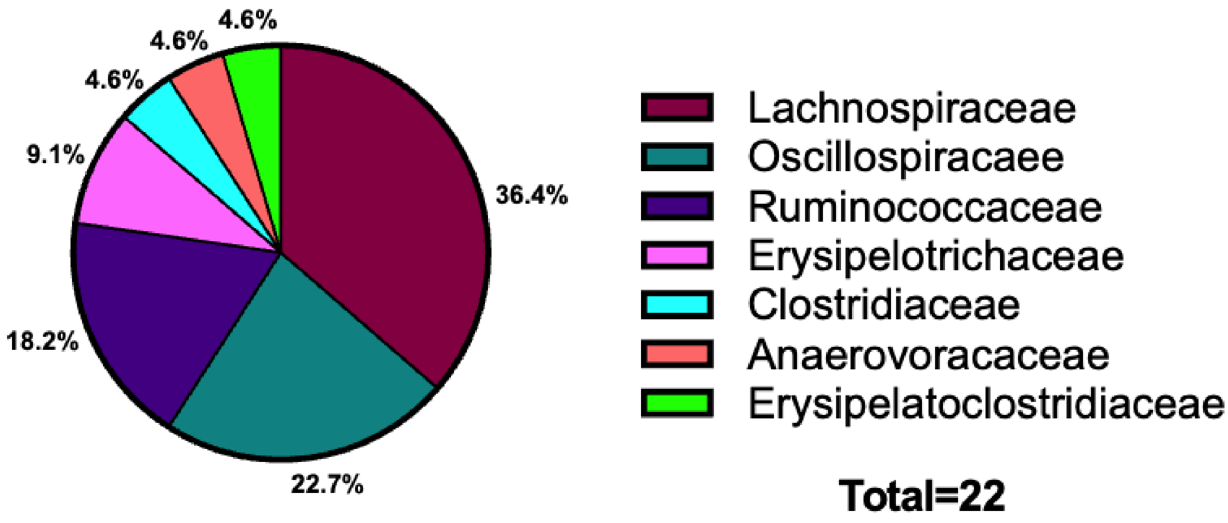


**Figure S2. Family-level distribution of cultivated ASVs.** Pie chart showing proportional family-level representation among the 22 unique ASVs cultivated on PYE6S and identified by 16S rRNA gene sequencing (total n = 22).

**Table S1.** Significant known metabolites from Volcano Plot in the positive and negative mode data sets. Metabolites in ALL CAPS are level 1 ID while lower case indicates level 3 IDs (less confident).

| Metabolite (Positive Mode) | FC | p.value |
|---|---|---|
| BUTYROBETAINE_146.1176-1.28 | 36.25 | 4.3745E-07 |
| 1,3-DIAMINOPROPANE_75.0917-0.59 | 9.42 | 0.0052908 |
| N-ACETYL-HEXOSAMINE_244.0791-0.81 | 7.47 | 2.1065E-09 |
| Kynurenic Acid_190.05-7.58 | 7.36 | 0.0073266 |
| D-Ribose_173.0421-0.77 | 6.49 | 1.6621E-09 |
| L-CYSTATHIONINE_223.0737-0.86 | 4.86 | 0.00020051 |
| GUANINE_152.0567-1.48 | 4.81 | 1.5458E-07 |
| Leu Pro_229.1546-7.18 | 3.80 | 2.2829E-05 |
| 2-hydroxyglutarate-water_131.034-1.51 | 3.75 | 1.0177E-06 |
| Glycerophosphocholine_258.109-0.76 | 3.52 | 0.0001874 |
| 4-HYDROXY-L-PHENYLGLYCINE_168.0652-0.81 | 3.51 | 1.8017E-06 |
| HEXOSE-6-PHOSPHATE_261.0368-0.71 | 3.42 | 0.003576 |
| ALPHA-AMINOADIPATE/N-METHYL-L-GLUTAMATE_162.0762-1 | 3.20 | 0.00030602 |
| Glycyl-L-leucine_189.1233-6.72 | 2.79 | 1.6905E-05 |
| Leu Pro_229.1545-7.38 | 2.77 | 0.00037724 |
| CARNOSINE_227.1135-0.7 | 2.62 | 4.0828E-06 |
| Dihydroxyquinoine_162.0549-7.97 | 2.48 | 0.0018949 |
| N-ACETYL-L-ASPARTIC ACID_176.0553-1.44 | 2.38 | 0.0018521 |
| L-METHIONINE_150.0584-1.39 | 2.36 | 0.00291 |
| NICOTINAMIDE_123.0552-1.6 | 2.35 | 0.00094456 |
| D-GLYCERIC ACID_129.0158-0.84 | 2.24 | 0.0092548 |
| 4-HYDROXY-2-QUINOLINECARBOXYLIC ACID_190.0501-7.19 | 2.23 | 0.00093408 |
| NICOTINATE_124.0395-1.37 | 2.15 | 9.6513E-05 |
| 2-AMINOPHENOL_110.0602-1.81 | 2.05 | 4.5585E-06 |
| ALDO/KETO-HEXOSE_203.0524-0.7 | 2.03 | 0.0082278 |
| DL-5-HYDROXYLYSINE_163.1075-0.61 | 0.49 | 0.010184 |
| 5-HYDROXY-L-TRYPTOPHAN_221.0921-6.82 | 0.47 | 0.0049216 |
| C5-SUGAR ALCOHOL_175.0584-0.74 | 0.46 | 0.021124 |
| Isovalerylcarnitine_246.1699-8.1 | 0.46 | 4.4384E-05 |
| L-ARGININE_175.119-0.81 | 0.46 | 0.077787 |
| QUINATE_193.0706-0.93 | 0.43 | 0.00031232 |

| | | |
|---|---|---|
| Pyroglutamic Acid_130.0499-1.78 | 0.43 | 0.00068947 |
| Monomethyl phthalate MMP_181.0494-9.21 | 0.42 | 0.00010257 |
| L-CYSTEINE_122.0269-0.75 | 0.38 | 8.7914E-07 |
| CAFFEATE_181.0495-7.67 | 0.34 | 7.6401E-06 |
| CYTIDINE_244.0926-1.69 | 0.33 | 0.007096 |
| L-Cystine_241.0308-0.69 | 0.33 | 2.4856E-06 |
| 4-GUANIDINOBUTANOATE_146.0924-1.59 | 0.32 | 0.0014047 |
| QUINATE_215.0524-0.91 | 0.32 | 2.1591E-05 |
| Choline_104.107-0.82 | 0.31 | 7.9093E-09 |
| D-GALACTOSAMINE_202.0684-0.67 | 0.28 | 2.1742E-06 |
| D-GLUCURONIC ACID/D-GLUCURONOLACTONE/D-GALACTURONIC ACID_217.0317-0.76 | 0.28 | 0.053716 |
| 3-SULFINO-L-ALANINE_154.0169-0.71 | 0.28 | 0.0028114 |
| Mannitol/Sorbitol_183.0863-0.73 | 0.27 | 0.0012396 |
| ADENINE_136.0618-1.48 | 0.26 | 0.00059579 |
| L-CARNITINE_162.1124-0.94 | 0.25 | 2.6414E-08 |
| 6C-SUGAR ALCOHOL_205.0683-0.73 | 0.24 | 0.0027511 |
| GLUCONIC ACID/D-GULONIC ACID GAMA-LACTONE_219.0472-0.73 | 0.22 | 6.7541E-08 |
| ALLANTOIN_159.0513-0.81 | 0.21 | 0.0037858 |
| 5-HYDROXYINDOLEACETATE_192.0656-7.43 | 0.21 | 0.00040112 |
| URATE_169.0356-1.79 | 0.21 | 5.1186E-06 |
| QUINATE_193.0705-0.83 | 0.14 | 4.6174E-11 |
| ORNITHINE_133.0971-0.62 | 0.13 | 0.011846 |
| 5-HYDROXYINDOLEACETATE_192.0656-7.57 | 0.11 | 4.856E-05 |
| LactoseK_381.0788-0.7 | 0.10 | 0.08509 |
| QUINATE_215.052-0.83 | 0.10 | 1.4514E-07 |
| D-GLUCOSAMINE 6-PHOSPHATE_260.0527-0.7 | 0.08 | 0.00032982 |
| Ibuprofen_207.138-8.11 | 0.07 | 1.0779E-10 |
| Lactose_343.123-0.72 | 0.06 | 0.015491 |
| D-RAFFINOSE_527.1578-1.09 | 0.05 | 0.001906 |
| HEXOSE-DISACCHARIDE_365.1049-0.7 | 0.04 | 0.0038421 |
| D-RAFFINOSE_527.1582-1.04 | 0.03 | 0.00013103 |
| D-RAFFINOSE_527.1582-0.98 | 0.01 | 8.8233E-05 |
| Trehalose/Sucrose_343.1232-0.94 | 0.01 | 0.030926 |

| **Metabolite (Negative Mode)** | **FC** | **p.value** |
|---|---|---|
| 3-Isopropylmalate_175.0611-1.16 | 21.94 | 0.05286 |
| L-METHIONINE_148.0434-1.57 | 12.88 | 0.0039892 |
| CHOLATE_407.2797-10.38 | 11.82 | 0.00019115 |
| N-ACETYL-D-MANNOSAMINE_256.0592-0.87 | 8.70 | 0.0039746 |
| N-ACETYL-D-MANNOSAMINE_256.0591-0.81 | 7.84 | 1.6487E-06 |
| N-ACETYL-HEXOSAMINE_220.0826-0.88 | 6.36 | 2.6862E-05 |
| GUANINE_150.0419-1.5 | 5.26 | 1.3976E-07 |
| N-ACETYL-HEXOSAMINE_220.0826-0.81 | 4.40 | 8.06E-08 |
| 3-METHYL-2-OXOVALERIC ACID_129.0558-6.68 | 3.59 | 0.00011117 |
| 2-hydroxyglutarate_147.0305-1.47 | 3.29 | 5.5878E-06 |
| GLUCOSE/FRUCTOSE_215.0328-0.72 | 3.19 | 4.3742E-06 |
| Glyceraldehyde_89.0239-0.75 | 3.19 | 9.6213E-08 |
| GLUCOSE/FRUCTOSE_217.0299-0.72 | 3.19 | 4.2093E-06 |
| PYRIDOXAMINE_203.0565-0.77 | 2.96 | 0.00013766 |
| N-ACETYL-L-ASPARTIC ACID_174.0407-1.54 | 2.85 | 0.0075053 |
| N-ACETYLNEURAMINATE_308.0985-0.87 | 2.79 | 0.0025943 |
| Chenodeoxycholic Acid CDCA_437.2906-12.38 | 2.65 | 0.0049953 |
| NICOTINATE_122.0247-1.38 | 2.27 | 0.00017014 |
| CITRULLINE_174.0883-0.78 | 2.27 | 0.006258 |
| N-ACETYL-L-ASPARTIC ACID_174.0409-1.46 | 2.23 | 0.0033233 |
| L-METHIONINE_148.0434-1.4 | 2.19 | 0.007222 |
| N-ACETYLNEURAMINATE_308.0986-0.8 | 2.09 | 0.021246 |
| ASCORBIC ACID-2H_173.0081-0.84 | 2.06 | 0.00057812 |
| CMP_322.0438-1.02 | 2.03 | 0.0085 |
| 4-COUMARATE_163.0399-8.23 | 0.46 | 0.050298 |
| 4-oxoproline_128.0353-1.79 | 0.44 | 0.0035534 |
| L-CYSTINE_239.0162-0.69 | 0.40 | 0.00016229 |
| 4-GUANIDINOBUTANOATE_144.0779-1.6 | 0.36 | 0.0043947 |
| GLUCONIC ACID/D-GULONIC ACID GAMA-LACTONE_195.0509-0.74 | 0.35 | 2.8793E-05 |
| ADENINE_134.0474-1.5 | 0.33 | 0.0019808 |
| N-AMIDINO-L-ASPARTATE_174.0501-0.93 | 0.32 | 1.2782E-05 |
| Mannitol_181.0712-0.74 | 0.31 | 0.00061463 |
| L-Cysteine-S-Sulfate_199.9692-0.75 | 0.30 | 0.00036535 |
| 2-AMINOETHYL DIHYDROGEN PHOSPHATE_140.0102-0.87 | 0.26 | 0.0010876 |
| ALLANTOIN_157.0372-0.81 | 0.24 | 0.0034947 |

| | | |
|---|---|---|
| URATE_167.0209-1.81 | 0.21 | 3.4713E-06 |
| D-SACCHARIC ACID_209.0304-0.75 | 0.21 | 0.00049624 |
| Nicotinamide ribotide_379.0504-0.96 | 0.17 | 0.0031141 |
| QUINATE_191.0561-0.94 | 0.15 | 1.3038E-05 |
| QUINATE_191.0554-0.83 | 0.14 | 1.3134E-11 |
| Vanilloylglycine_224.0564-7.64 | 0.12 | 6.6224E-10 |
| D-GLUCOSAMINE 6-PHOSPHATE_258.0388-0.74 | 0.09 | 0.0011957 |
| SHIKIMATE_173.0455-1.08 | 0.08 | 4.7977E-12 |
| D-RAFFINOSE_503.1614-1.03 | 0.06 | 0.0014471 |
| D-RAFFINOSE_503.1613-0.89 | 0.05 | 0.00077307 |
| Ferulic acid sulfate_273.0076-8.17 | 0.03 | 0.00018809 |
| STACHYOSE_665.2138-0.96 | 0.02 | 0.024194 |
